# Single-nucleus proteomics reveals proteins regulating LPS-induced nucleocytoplasmic transport in macrophages

**DOI:** 10.1101/2024.06.17.599449

**Authors:** Jason Derks, Tobias Jonson, Andrew Leduc, Mahmoud-Reza Rafiee, Saad Khan, Luke Khoury, Nikolai Slavov

## Abstract

**Background:** Variation in the proteome configuration of unstimulated cells may be associated with differences in how individual cells respond to stimulation. Multiplexed single-cell proteomics by liquid chromatography mass spectrometry can uncover these associations at high depth and throughput.

**Results:** Here we develop an approach that links proteomic and phenotypic variation across individual cells and nuclei to identify potential regulators of biological processes for targeted validation. We apply this approach to identify regulators of nucleocytoplasmic protein transport in macrophages stimulated with lipopolysaccharide (LPS). We quantify the proteomes of 3,412 individual nuclei across the dynamic response to LPS and associated proteomic variation with differences in transport. Within minutes of stimulation, protein transport in individual nuclei correlates strongly with the abundance of known protein transport regulators, revealing how natural protein variation relates to cellular responses. Simple biophysical constraints, including nuclear pore abundance, partially explain the variation in LPS-induced nucleocytoplasmic transport. We select 16 new potential regulators of protein transport to test by targeted knockdown. The knockdown phenotypes confirm the inferences derived from natural protein and functional variation of single nuclei.

**Conclusions:** By linking proteomic and functional variation among individual nuclei, we identify proteins associated with LPS-induced nucleocytoplasmic transport and prioritize potential regulators for functional validation. This approach should be broadly applicable to discovering protein regulators and enhancing the functional interpretation of single-cell omics data.

## Background

Single-cell omics methods have rapidly scaled up^1,2^ and enabled the investigation of cellular heterogeneity and the creation of cell atlases^3^. However, the functional interpretation of such omics data has lagged behind^4–7^. To investigate the link between proteomic and functional variability across single-cells, we sought to develop an approach that associates the pre-existing differences in proteome configurations to their correspondingly variable cellular responses; such an approach may enable the inference of novel regulatory associations^8,9^. In particular, we aimed to investigate how pre-existing proteomic variability influences, and therefore explains the variability of, lipopolysaccharide (LPS)-induced transport of proteins to and from the nucleus.

To quantify this variation between individual cells and individual nuclei, we built upon the conceptual and technological advances in single-cell mass spectrometry proteomics^10–24^. Specifically, we used the framework of multiplexed data-independent acquisition (plexDIA) to implement the isotopologous carrier suggestion^25,26^, which generalizes the concept of isobaric carrier^27,28^ to plex-DIA. Isotopologous carriers have shown promise in some applications^15,29–31^, and here we used them to enable the first proteomic analysis of individual organelles isolated from human cells. These methodological advances enabled global exploration of protein transport within individual cells, which for decades has been studied by imaging fluorescent proteins^32–36^. Our approach allows for more comprehensive initial discovery, which can expand previous observations that pro-inflammatory stimulation leads to heterogeneous nuclear import of transcription factors^37,38^. These discoveries can later be examined using directed perturbations or higher temporal resolution using fluorescent imaging^39,40^.

Here, we demonstrate a generalizable approach that enables the inference of functional regulators from (sub-)single-cell proteomics data. Specifically, we identified proteins regulating subcellular transport, including the differential contributions of the subunits of the nuclear pore complex. Our inferences were derived from natural protein variation across single cells and nuclei. Subsequent validation of these results by targeted knockdown experiments provide direct evidence for the functional relevance of our inferential approach.

## Results

### Protein transport in bulk macrophage populations

To study nuclear spatiotemporal changes in bulk cell populations, six biological replicates were generated using a model system of macrophage-like cells derived from THP-1 monocytes treated with phorbol 12-myristate 13-acetate (PMA). Nuclei were extracted with mild detergent using methods adapted from an established workflow^41^. This physical isolation produced nuclear enrichment as demonstrated by a 5 - 25-fold depletion of proteins from non-nuclear cellular compartments from the nuclei, though variability was observed between biological replicatesAdditional file 1: Fig. S1. A separate experiment was performed to evaluate nuclear membrane integrity using fluorescent dextran of variable masses, though the results were inconclusive Additional file 1: Fig. S2. To mitigate influences of permeability and enrichment variability on downstream interpretation, six biological replicates were analyzed.

Having validated the depletion of non-nuclear proteins from the isolated nuclear fractions, we sought to quantify the LPS-induced dynamics of protein transport. To this end, we quantified differential protein abundances for bulk populations of nuclei and whole-cells using MS-EmpiRe^42^. The abundance changes in the whole cells reflect protein synthesis, degradation, and secretion; changes at the nuclear level also include nucleocytoplasmic transport. As expected, changes mediated by protein synthesis and degradation are slow compared to the kinetics of nucleocytoplasmic transport. Indeed, 60 minutes after LPS-stimulation, the changes are dominated by transport with 15.7% of nuclear proteome exhibiting differential abundance compared to only 2.3% of the whole-cell proteome, Fig. 1a. This difference is even more pronounced for earlier time points where changes affect less than 0.1% of the whole-cell proteome 30 minutes after LPS stimulation, but 9.3% of the nuclear proteome.

**Figure 1.**
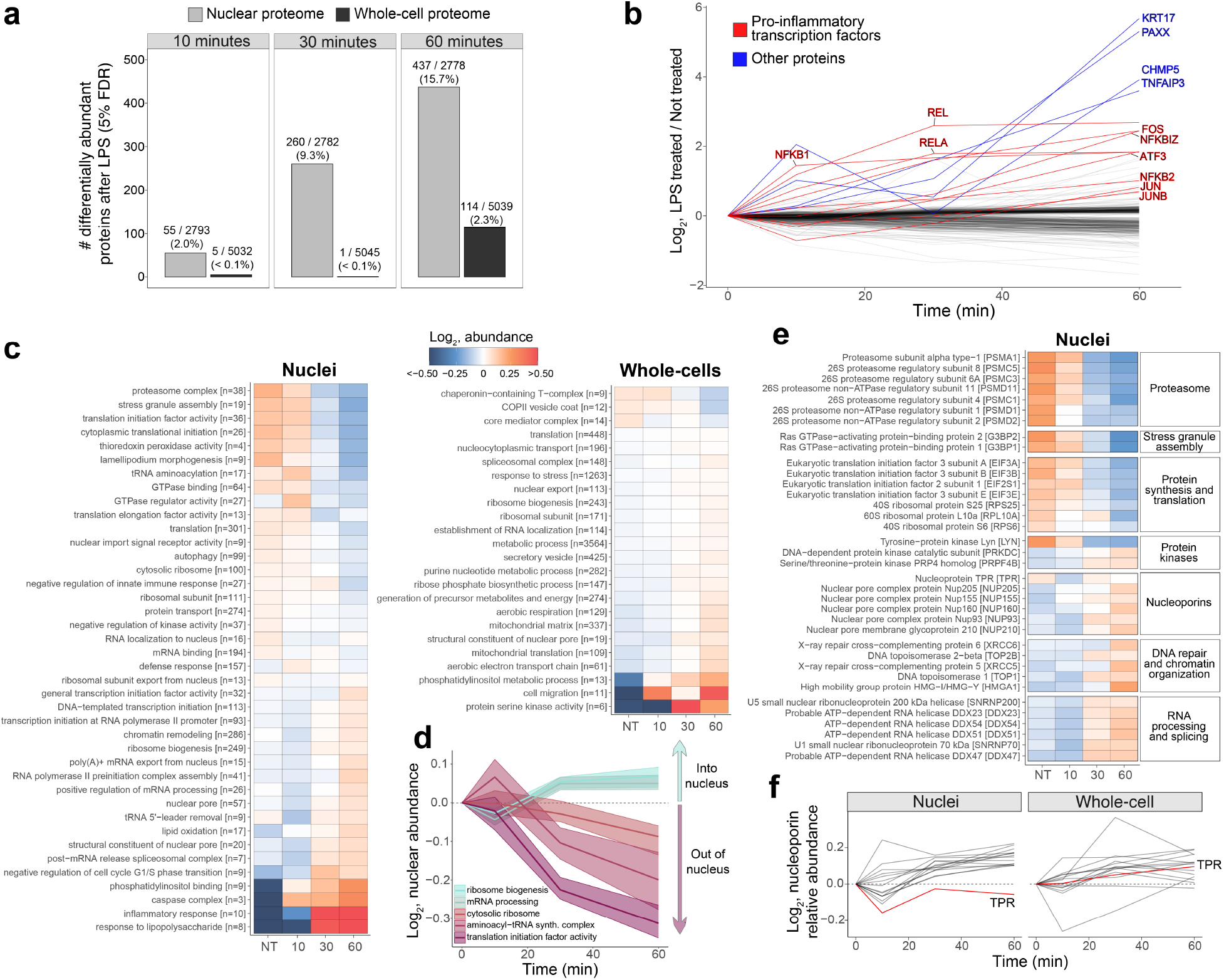
Protein dynamics in whole cells and nuclei reveal coordinated spatiotemporal control of biological processes in bulk populations of macrophage-like cells. **a** Number of differentially abundant proteins (5% FDR) in nuclear (gray) and whole-cell (black) proteomes after 10, 30, or 60 minute LPS treatments. **b** Time-series data over the course of the LPS-treatment for differentially abundant proteins in the nucleus; pro-inflammatory transcription factors are labeled and highlighted in red, and other proteins with *>*4-fold change are highlighted in blue. **c** Protein set enrichment analysis of nuclei and whole-cell bulk samples for not-treated (NT), 10 minute, 30 minute, and 60 minute LPS treated samples. Brackets correspond to the number of proteins included in computing the enrichment of Gene Ontology terms. **d** Gene Ontology terms associated with mediating gene-expression, which significantly change in nuclear abundance in response to LPS, are plotted. The 95% confidence intervals are derived from the change of all proteins which correspond to the respective Gene Ontology term. **e** Proteins were grouped thematically and plotted to show their change in nuclear abundance; all proteins were differentially abundant at 5% FDR in at least one time point. **f** Nucleoporins which were differentially abundant in the nucleus in at least one time point are plotted to display their change in nuclear and whole-cell abundance; TPR is highlighted in red.

Given that hundreds of proteins without associations to innate immunity significantly changed their nuclear abundance in response to LPS, we aimed to evaluate the temporal continuity of each protein to further assess confidence in these findings. As a quantifiable continuity metric, we calculated a ranked version of von Neumann’s ratio (RVN)^43^ for each protein as shown in Additional file 1: Fig. S3a. Proteins found to be differentially abundant at 5% false discovery rate (FDR) for nuclei and whole-cells experienced mostly monotonic changes that exceeded a simulated null, Additional file 1: Fig. S3b,c, which supports the validity of the observed spatiotemporal changes.

### Mass dependence of nucleocytoplasmic transport

The kinetics of nucleocytoplasmic transport is mass-dependent, and we sought to investigate this dependence in our data. There are two mechanisms by which transport occurs: passive diffusion and active transport^44,45^. Molecules with masses less than 40 kDa have been reported to passively diffuse through nuclear pores with kinetics that are mass-dependent^46^. Kinetics of active transport are also dependent on the size of the cargo, but to a lesser degree, as the energy barrier is reduced through interactions with chaperones^44,47^. We investigated this relationship in our data and found that protein transport was indeed negatively correlated with protein mass, as shown in Additional file 1: Fig. S4. While this analysis cannot conclude whether these trends reflect passive diffusion or simply mass-dependence in active transport, the results are consistent with more recent findings of passive diffusion occurring without a precise size threshold^48^ and corroborate the mass-dependence of protein transport kinetics globally in the natural response of macrophages to LPS.

### Spatiotemporal control of inflammatory response in bulk cell populations

While pro-inflammatory transcription factors are known to be imported to the nucleus in response to LPS^49–51^, we identify hundreds of additional proteins significantly changing in nuclear abundance, many of which have no prior association with LPS-response. Indeed, global subcellular proteomic rearrangement has been previously observed in response to LPS^51^. As expected, transcription factors NF-*κ*B1, REL, RELA, and FOS increase in nuclear abundance by approximately 4-fold within 10-60 minutes of LPS-treatment^33,36,37,52^, as shown in Fig. 1b. Interestingly, hundreds of additional proteins experience monotonic or generally continuous changes in nuclear abundance; one of these proteins, PAXX––a protein required for non-homologous end joining (NHEJ) DNA repair^53,54^, reached a 40-fold increase within 60 minutes of LPS-stimulation. This protein acts as a scaffold for the accumulation of XRCC5 and XRCC6 to facilitate NHEJ DNA repair^55^. Likewise, XRCC5 and XRCC6 also increased significantly in the nucleus by 60 minutes, Additional file 1: Fig. S5.

To characterize the effects of protein synthesis, degradation, and secretion, we computed protein set enrichment on whole-cell proteomes; similar analysis of nuclear proteomes would additionally reflect nucleocytoplasmic transport, Fig. 1c. LPS-stimulation induced whole-cell pro-teome enrichment of Gene Ontology (GO) terms associated with gene-expression, such as “translation,” “spliceosomal complex,” and “ribosome biogenesis.” Interestingly, changes to the nuclear proteome revealed spatiotemporal changes consistent with this whole-cell enrichment; specifically GO terms whose sites of action are in the nucleus, such as “general transcription initiation factor activity” and “ribosome biogenesis” increased in nuclear abundance, while GO terms whose sites of action are outside of the nucleus, such as “cytosolic ribosome” decreased in nuclear abundance. These changes which reflect coordinated spatiotemporal rearrangement of proteins involved in mediating gene-expression are shown for five GO terms in Fig. 1d. Together, the whole-cell and nuclear proteomic data provide complementary evidence consistent with increased gene-expression in response to LPS, as observed through increases in the absolute abundances of and the spatiotemporal rearrangement of proteins involved in this process.

To more closely investigate the concordance of changes in nuclear biological processes, differentially abundant proteins were grouped thematically by function and plotted in Fig. 1e. Proteins associated with DNA repair, RNA processing, and nucleoporins were generally found to increase in nuclear abundance following LPS stimulation. Intriguingly, of all nucleoporins which changed significantly in nuclear abundance, only TPR––a negative regulator of nuclear pore complex (NPC) assembly^56^––decreased, Fig. 1f. Whole-cell abundances of the same nucleoporins were generally found to increase, including TPR. The observed whole-cell and subcellular changes are consistent with LPS-induced upregulation of NPC assembly.

### Benchmarking protein quantification of single nuclei

Using natural variation across single-cells to identify regulators of protein transport requires accurate quantification of many proteins in single nuclei. To accomplish this, we used highly parallel sample preparation by nPOP^16,57^ and the plexDIA framework combined with an isotopologous carrier. This methodology is analogous to the isobaric carrier introduced by SCoPE-MS^12,27,58^, with the exception that the mass tags are isotopologous and thus separable at MS1 and MS2 levels. As previously suggested and demonstrated with plexDIA, this framework uses the chromatographic coelution of peptides from different samples to increase data-completeness^25,26^. Thus, protein identification across all samples can be enhanced by including a highly abundant sample, e.g., an isotopologous carrier, in parallel to single-cells or single-organelles. This benefit was evident with the introduction of plexDIA^25^ and has been utilized for single-cell analysis^15,31^. However, complex isotopologous carriers might increase interferences and thus decrease quantitative accuracy. To evaluate these trade-offs and find an optimal carrier level for acquiring single-nucleus data, we created and acquired data from a mixed-species spike-in of *H. sapiens* nuclei and *S. cerevisiae* run with 0x, 1x, 5x, 10x, 25x, or 50x carrier amounts, as shown in Additional file 1: Fig. S6a. We found that intersected protein-level quantitation was similar but slightly decreasing at higher carrier-levels, Additional file 1: Fig. S6b. Thus, the potential for interference by larger carriers has little effect on quantitative accuracy in our experiments, which is likely due to the lower proteomic complexity of nuclei as compared to whole cells.

To improve quantitative accuracy, the carrier channel was used as a reference to estimate quantitative compression and remove precursors which consistently show high levels of compression, similar to previous carrier based filters^12^. We applied this filter at various levels and found that removing precursors in single nuclei-level inputs which were systematically compressed *>* 6-fold relative to the carrier struck an optimal balance between retained coverage and accuracy, Additional file 1: Fig. S6c-e. Using this strategy and filtering level, we compared carrier levels without intersecting proteins to show the complete coverage and quantitative accuracy for each, Additional file 1: Fig. S6f. While intersected quantitative accuracy was similar across carrier levels, a decrease in accuracy was observed at increasing carrier levels on non-intersected data. This decrease in accuracy corresponds to the quantification of less abundant proteins in single-nucleus channels. Given these results, 25x and 50x nuclear proteome carriers were used to acquire single-nucleus data.

### Variability in LPS-induced nucleocytoplasmic transport across single nuclei

Next, we sought to infer potential regulators of nucleocytoplasmic transport from the natural variability of macrophage proteomes prior to the LPS stimulation and its regulatory impact on the LPS response. Using SCoPE-DIA, we acquired a dataset consisting of 3,412 nuclei across four biological replicates, with a median of 1,366 proteins quantified per nucleus, Fig. 2a. After filtering to remove nuclei and precursors with poor quantitation (Additional file 1: Fig. S7a-b), and nuclei with insufficient nuclear enrichment (Additional file 1: Fig. S7c-d), we retained 2,997 nuclei with a median of 1,287 proteins per nucleus to be used in downstream analyses, Fig. 2a.

**Figure 2.**
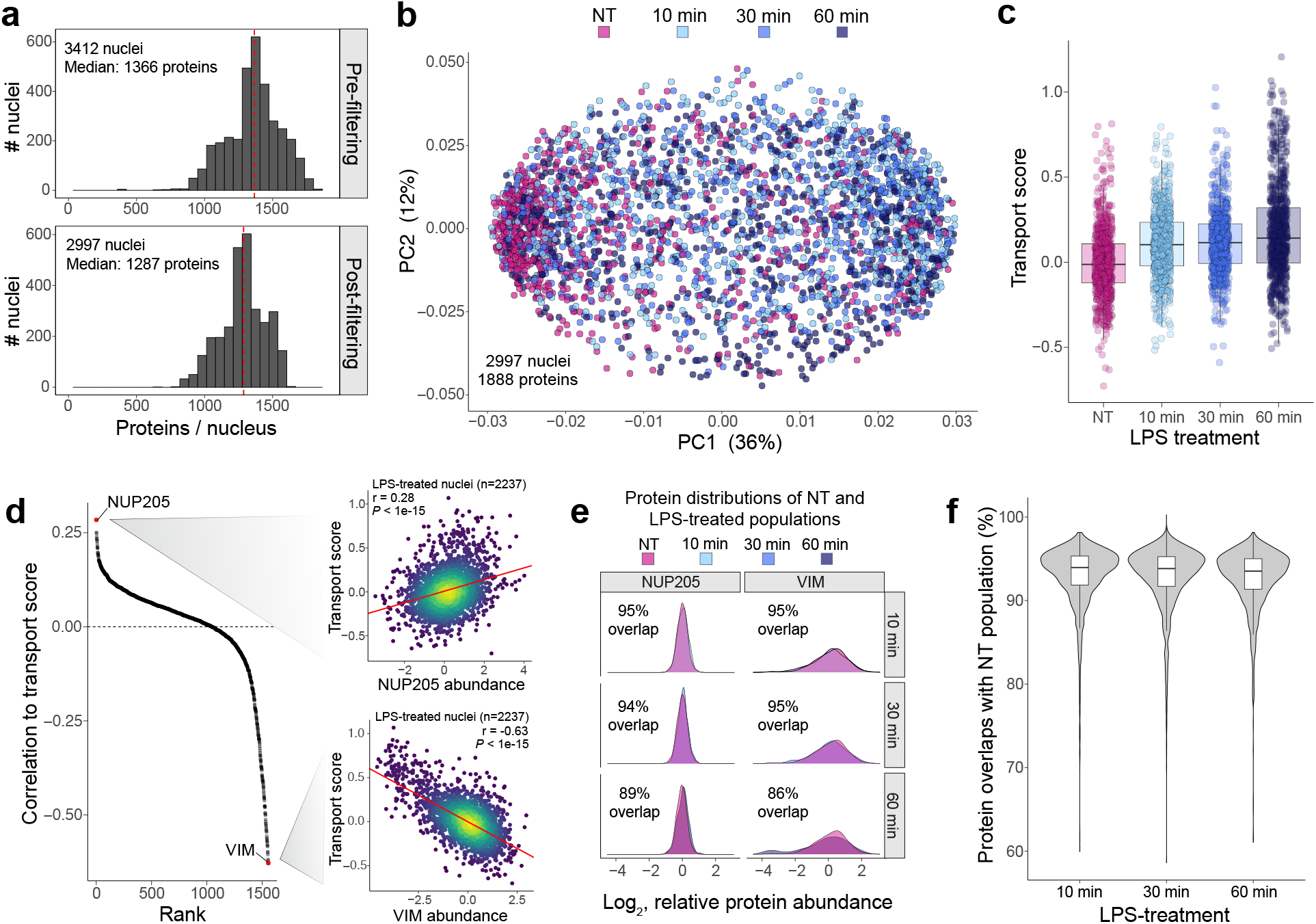
Associating single-nucleus proteomic variability to heterogeneous LPS-induced nucleocytoplasmic protein transport. **a** Liquid chromatography–tandem mass spectrometry (LC-MS/MS) single-nucleus proteomics data was acquired with SCoPE-DIA for 3412 nuclei, quantifying a median of 1366 proteins per nucleus. Single nuclei which were unsatisfactorily prepared, acquired, or depleted of non-nuclear proteins were removed, yielding 2997 nuclei with a median of 1287 proteins per nucleus post-filtering; these nuclei were used for downstream analyses. **b** Weighted principal component analysis (PCA) produced partial separation between LPS-treated (10, 30, and 60 minute) nuclei and not-treated (NT) nuclei. **c** Distribution of transport scores from single nuclei. Macrophages treated with LPS for longer durations yielded nuclei which had undergone more protein transport. **d** Rank sorted correlations between the transport score of each nucleus and its proteome. The underlying data used to compute these associations is shown for two proteins, NUP205 and VIM. **e** Distributions of relative protein abundances of NUP205 and VIM in the NT and LPS-treated populations of single nuclei. The overlap quantifies the proportion of commonality between the untreated and the treated populations. High values suggest pre-existing variability is much larger than LPS-induced changes. **f** Overlap coefficients were computed for all proteins between NT and LPS-treated populations of single nuclei and the distributions were plotted.

Projecting the single-nuclei proteomes onto their principal components produced a partial separation between NT and LPS-treated populations, Fig. 2b. Aside from measurement error, three compounding sources of variability may result in this observed partial separation: pre-existing variability of protein abundances at the whole-cell level, pre-existing variability in subcellular protein localization, and variability in nucleocytoplasmic transport in response to LPS. We sought to overcome the first two sources of variability and investigate the latter, heterogeneity of nucleocyto-plasmic transport. This presents a challenge as the pre-existing protein variability is large relative to the magnitude of protein transport induced by a short LPS exposure.

To explore the variability of LPS-induced nucleocytoplasmic transport, we derived a metric aiming to quantify the amount of protein transport experienced by each nucleus; however, given that we did not measure proteomes of the same nuclei before and after stimulation, our estimate cannot directly quantify protein transport. As a substitute, we derived a ‘transport score’ metric, which quantifies the deviation of single LPS-treated nuclear proteomes from the proteome distribution of untreated single nuclei. To mitigate noise, the bulk data was used to differentially apply weights based on transporting proteins, as illustrated in Additional file 1: Fig. S8. As expected, the resulting distributions of transport scores for single nuclei indicate increased nucleocytoplasmic transport for longer time-periods of LPS treatment, Fig. 2c.

To link proteomic variability to functional variability in protein transport, we computed correlations of relative protein abundances to transport scores for all LPS-treated single nuclei, Fig. 2d; we highlight two proteins, NUP205 (*r* = 0.28) and VIM (*r* = *−*0.63), which are among the most positively and negatively correlated proteins to protein transport. Results from protein-set-enrichment analysis that was performed on this ordered vector indicate associations to cell division, cell adhesion, and other processes shown in Additional file 1: Fig. S9. To explore whether these associations are driven by pre-existing proteomic variability or LPS-induced changes, we compared the protein abundance distributions before and after LPS treatment, Fig. 2e. The distributions were highly similar between conditions, suggesting LPS-induced changes were small relative to the natural variation across untreated nuclei. We evaluated this overlap systematically for all protein between NT and LPS treated conditions and plotted the results for all protein overlap coefficients in Fig. 2f, which likewise indicate highly similar distributions for protein abundances between NT and LPS treated conditions. Therefore, the associations are dominated by pre-existing variation across proteomic configurations; this initial variability likely influences the amount LPS-induced protein transport each cell undergoes. Thus, associations between single-nucleus protein abundances and transport scores may be used to identify proteins whose abundance affected nucleocytoplasmic protein transport.

### More nuclear pore complexes, more transport

Nearly all nucleoporins were positively correlated to nucleocytoplasmic protein transport, Fig. 3a, suggesting nuclei with more nuclear pore complexes (NPCs) experience more transport. The only exception was TPR, which has been reported to negatively regulate NPC assembly, as validated through knockdowns of TPR yielding cells with more assembled NPCs^56^, as well as in patient fibroblasts with reduced TPR abundances^59^. Interestingly, our data support these previously observed results. To more clearly investigate the relationship between NPC abundance and protein transport, we collapsed all nucleoporin abundances to the complex-level, and correlated these NPC abundances to transport scores in LPS-treated nuclei (n = 2,237) and found a relatively strong and highly significant association (*r* = 0.48, *P* < 10^−15^), as shown in Fig. 3b. Indeed, this suggests a highly significant association between NPC abundance and nucleocytoplasmic transport, which has been observed in the context of aging cardiomyocytes^60^.

**Figure 3.**
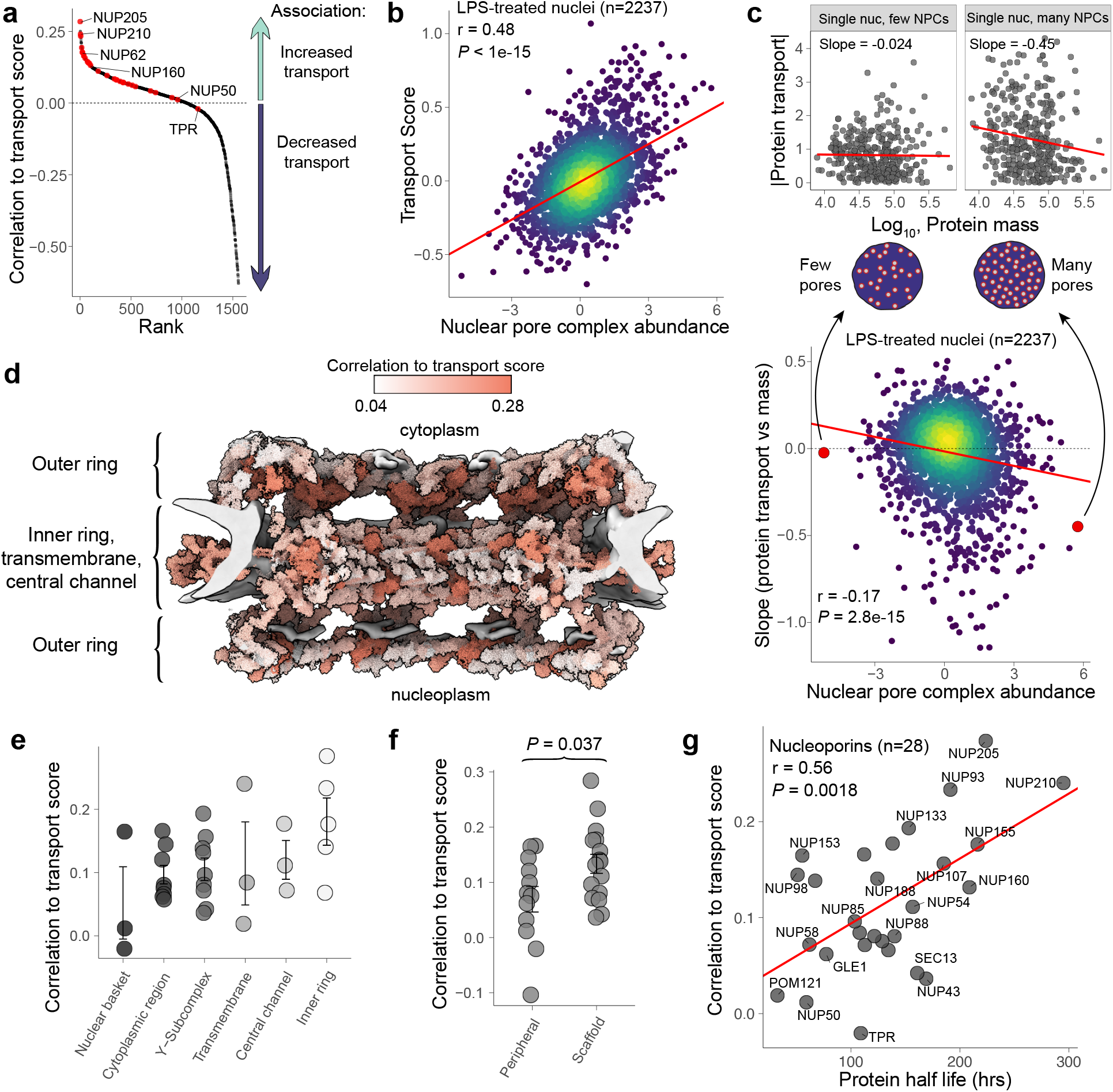
Nucleoporin abundances and their associations to LPS-induced nucleocytoplasmic transport in single nuclei. **a** Nucleoporins are variably associated with LPS-induced nucleocytoplasmic protein transport. The x-axis is ordered by protein rank. **b** Single nuclei with more nuclear pore complexes experience more LPS-induced nucle-ocytoplasmic protein transport (*P<* 10*^−^*^15^, *r* = 0.48). **c** For each single LPS-treated nucleus, a slope of the absolute value of differentially abundant proteins vs protein mass was computed to quantify mass-dependence of transport for each single nucleus; single nuclei which had the least and greatest abundance of nuclear pore complexes are shown at the top of the panel. The resulting slopes were plotted against their respective nuclear pore complex abundances, as shown at the bottom of the panel. The data suggest protein transport becomes increasingly mass-dependent, favoring transport of smaller proteins, when nuclear pore complex abundances are high (*P<* 10*^−^*^14^, *r* = *−*0.17). **d** Predicted human nuclear pore complex from Mosalaganti *et al.*, with nucleoporins colored by their correlations to transport score. **e** Distribution of each nucleoporin’s correlation to transport score grouped by their localization inside the NPC. **f** Distribution of each nucleoporin’s correlation to transport score grouped by whether it is a peripheral or scaffold protein in the NPC. The statistical significance between these two groups was tested using a two-sided Student’s *t*-test (*P* = 0.037). **g** Nucleoporins’ correlations to transport are associated with their respective half-lives (half-life data from Mathieson *et al.*).

Having uncovered this association between transport and NPC abundance in single nuclei, we sought to additionally investigate whether the mass-dependence of nucleocytoplasmic transport also changes as the number of NPCs varies across single nuclei; specifically, the nuclear envelope should become more permeable to passive diffusion at higher pore densities. To test this hypothesis, we first quantified the mass dependence of protein transport for each nucleus by regressing the molecular masses of proteins on their absolute deviations from the NT-population of single nuclei––a substitute for measuring protein-specific transport. The slopes of these regressions were plotted against corresponding NPC abundances in single nuclei, Fig. 3c. The results revealed a weak association across all 2,237 LPS-treated nuclei (*r* = *−*0.17) with high statistical significance (*P <* 10*^−^*^14^). This association indicates that nucleocytoplasmic transport of smaller proteins increasingly outpaced transport of larger proteins, at least in part as a function of NPC abundance, Fig. 3c. Given that passive diffusion is highly mass-dependent, it may suggest that single nuclei with more NPCs experience more passive diffusion.

While the abundances of nearly all nucleoporin proteins correlate positively with transport scores, the magnitude of this correlation varies across nucleoporin proteins, as shown in Fig. 3a. To explore this variability, we investigated what might be associated with these differences. To visually inspect the structural dependence of the correlations, the predicted structure of the human nuclear pore complex from Mosalaganti *et al.*^61^ was colored according to each proteins’ correlation to the transport score, Fig. 3d. This was investigated more directly by grouping nucleoporins based on their localizations inside the nuclear pore complex, as shown in Fig. 3e, and according to whether the nucleoporins were peripheral or scaffold proteins of the NPC^62^, as shown in Fig. 3f. The abundances of scaffold proteins were significantly more associated with nucleocytoplasmic transport than peripheral proteins of the NPC (*P* = 0.037). This variation across nucleoporins is further supported by a significant correlation with protein half-lives derived from Mathieson *et al.*’s stable isotope labeling by amino acids in cell culture (SILAC) turnover data^62^ (*r* = 0.56, *P* = 0.0018), as shown in Fig. 3g, as scaffold nucleoporins have been reported to have longer half-lives than peripheral nucleoporins^62,63^. These results suggest the observed variability in each nucleoporins’ correlation to transport is significantly associated with the nucleoporin’s localization inside the NPC and their respective half-lives.

### Targeted perturbations validate the single-nucleus derived associations

Having used single-nucleus proteomics data to infer novel regulators of LPS-induced nucleocyto-plasmic protein transport, we aimed to directly test these potential enhancers and suppressors of nucleocytoplasmic transport through targeted perturbations. The highest ranking hypotheses from the single-nucleus proteomics analysis include proteins directly involved in transport such as nucleoporins and ribosomal export proteins such as MDN1^64^. However, many other proteins without functional annotations for protein transport or innate immunity are also highly (anti-)correlated to transport. Perhaps some of these proteins influence LPS-induced protein transport, and these functional associations present discovery opportunities. We sought to test these associations experimentally through directed perturbations using siRNA mediated knock downs.

To test potential transport regulators, we chose to knock down 16 proteins spanning the continuum shown in Fig. 4a and quantified the change in LPS-induced nucleocytoplasmic protein transport. The knockdowns were performed using siRNAs as shown in Fig. 4b. The decrease in the corresponding gene products was validated in whole-cell and nuclear bulk lysates, as shown in Additional file 1: Figs. S10 and S11 resulting in the depletion of target proteins approximately 2- to 1.2-fold. To quantify how each knockdown affected LPS-induced nucleocytoplasmic protein transport, slopes were calculated from fold-changes (LPS/NT) between the original bulk data and each siRNA condition, using the negative control (non-targeting) siRNA as a reference for each biological replicate, as shown in Fig. 4b. Results for individual biological replicates are shown in Additional file 1: Fig. S12. With this strategy, we aimed to quantify the global change in nucleo-cytoplasmic transport as a result of each knockdown.

**Figure 4.**
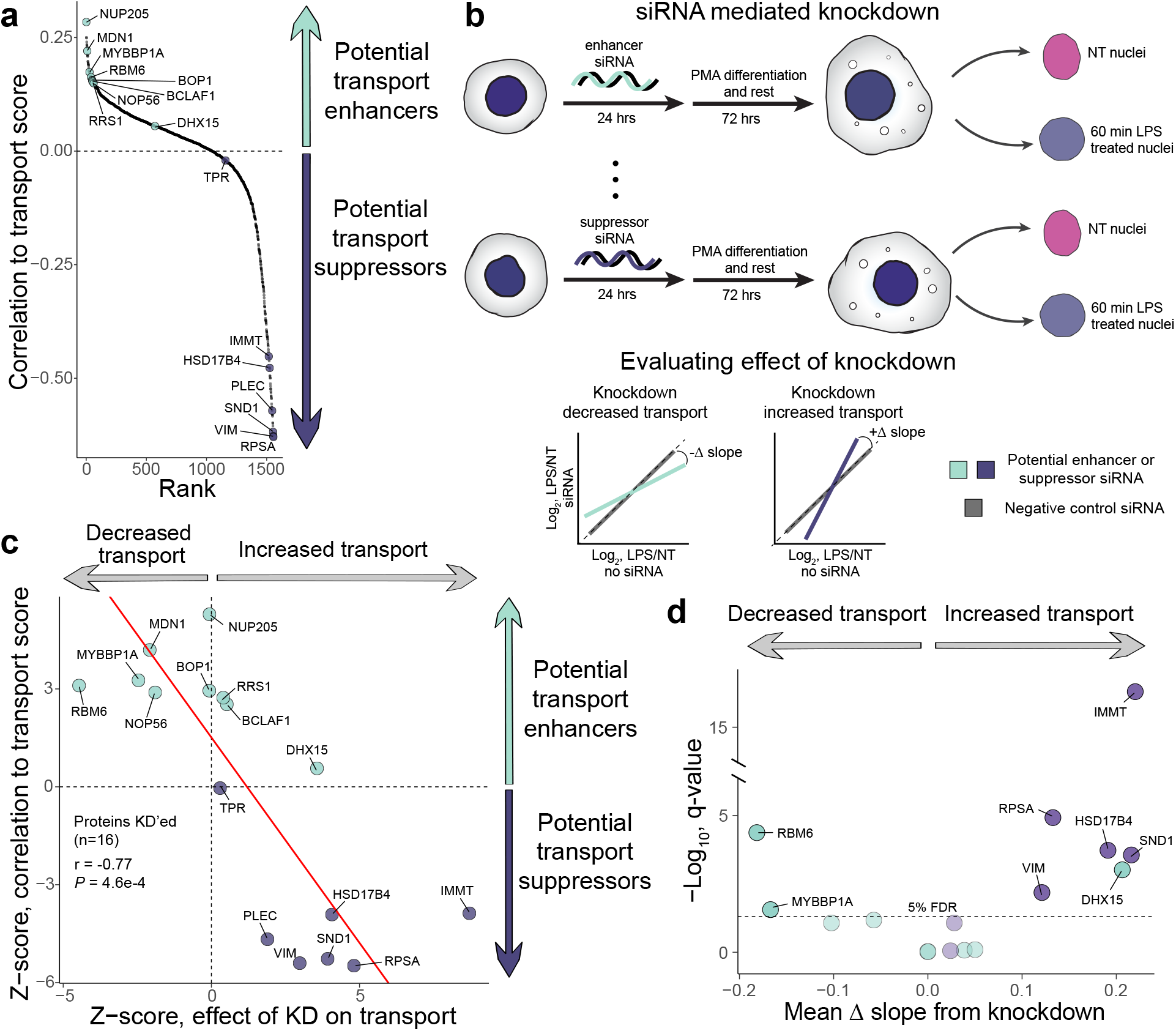
Genetic perturbations validate single-nucleus-derived protein associations. **a** Correlations of protein abundances to transport score from single-nucleus data. 16 proteins were selected to be knocked-down and colored with green or purple depending on whether it is associated with increased or decreased transport. **b** Workflow for siRNA-mediated knockdowns in THP-1 monocytes, followed by differentiation to macrophages. A negative control (non-targeting) siRNA was included in each biological replicate, and used to compute a change in transport as the result of each siRNA knockdown (KD); change in transport was calculated as the difference in slopes between the negative control siRNA and a gene-targeting siRNA. **c** Here we evaluate the predictive power of the single-nucleus findings through experimental genetic perturbations in bulk data. The biological replicates from the single-nucleus data were used to compute significance of correlations to transport scores, shown on the y-axis; the x-axis measures the significance of how the knockdown affected nucleocytoplasmic protein transport. The single-nucleus associationswere highly predictive of the functional effects in genetic perturbations (*P* = 0.00046, *r* = *−*0.77). **d** Significance of how individual gene knockdowns affect nucleocytoplasmic transport. Knockdowns of IMMT, RPSA, HSD17B4, SND1, DHX15, and VIM were found to globally increase LPS-induced nucleocytoplasmic protein transport; knock-downs of RBM6 and MYBBP1A were found to decrease transport.

Associating relative protein abundances to function in single cells may serve as a means of inferring regulatory potential. We sought to evaluate the predictive power of these associations by computing the significance of protein correlations to transport from the single-nuclei data and the significance of the change in transport from corresponding knockdowns, Fig. 4c. Indeed, as a result of the knockdowns, protein transport for 13/16 of the conditions was affected in the expected direction. Specifically, knockdowns of potential transport suppressors increased transport in 7/7 cases; knockdowns of potential transport enhancers decreased transport in 6/9 cases. The protein correlations to transport scores from the single-nucleus data were highly predictive of the functional effects of the knockdowns (*r* = *−*0.77, *P <* 5 *×* 10*^−^*^4^). The slopes, which were used for determining the change in LPS-induced nucleocytoplasmic transport, were computed from the 100 most differentially abundant proteins from the original bulk data. Various filtering levels were applied to examine the robustness of the findings, as shown in Additional file 1: Fig. S13, and all three levels showed similar trends.

Knockdowns of IMMT, RPSA, HSD17B4, SND1, DHX15, and VIM were found to significantly increase transport, whereas knockdowns of RBM6 and MYBBP1A were found to significantly reduce transport, Fig. 4. Therefore, 8/16 knockdowns significantly changed LPS-induced nucleocytoplasmic transport, 7/8 of which changed nucleocytoplasmic transport in the expected direction. The knockdown which most significantly affected transport was IMMT, a mitochondrial protein without prior association to innate immunity or nucleocytoplasmic protein transport. Whole-cell lysates of these knockdowns complementarily indicate a gradient of unpreparedness and primedness to respond to LPS and facilitate nucleocytoplasmic transport, as shown in Additional file 1: Fig. S14. These findings validate the potential of (sub-)single-cell proteomics to identify novel regulators of biological functions.

## Discussion

Identifying functional regulators of complex biological processes is an ongoing challenge in omics data interpretation. Here we demonstrate an interpretation approach that exploits the natural variability inherent in a population of single cells to empower inference of functional regulation. It is conceptually similar to the identification of functional regulators through perturbation-induced variability in bulk samples^65^ but instead utilizes the natural variability across individual cells. In our approach, we assumed that the initial proteomic states of single macrophages would explain the variability in LPS-induced nucleocytoplasmic protein transport. Operating under this principle, we quantified a metric (the transport score) from which we could evaluate the influence of resting proteomic variability on this response. We found that simple biophysical constraints, such as the quantity of nuclear pores, partially explain the variance in nucleocytoplasmic transport. Interestingly, the abundance of scaffold subunits is more strongly associated with transport rates than the abundance of peripheral subunits, which is consistent with modular and specialized structures of the nuclear pore complex^66^.

Beyond the role of nuclear pores, our analysis identified hundreds of additional proteins, many of which without annotated associations to innate immunity or protein transport, yet whose abundances were strongly correlated to the magnitude of nucleocytoplasmic protein transport experienced in single cells. We tested these protein-associations through genetic perturbations and found that the associations derived from single-nucleus data were, as a whole, highly predictive of their respective effects on LPS-induced nucleocytoplasmic transport. One consideration with these results is that the knockdowns produced relatively modest protein depletions. Yet, they had the expected result on the biological phenotype of interest: the rate of nucleocytoplasmic protein transport. The knockdowns changed the rate of protein transport to a degree and a direction predicted by the protein covariation across single nuclei, indicating strong functional validation of the single-nucleus proteomics results and approach.

A key enabling factor of our analysis is the ability to accurately and comprehensively quantify the proteomes of thousands of individual nuclei. Thus, the power of this approach will grow as single-cell proteomics becomes more accurate and higher throughput. Methods that increase throughput by shortening separation times^67^, using higher-plex tags^68^, and multiplexing samples in time^69^ will produce larger datasets that will further refine hypotheses derived from natural protein variation by allowing better discrimination between direct and indirect regulatory interactions^11^.

Future work will also benefit from extending the scope of protein measurements (such as quantifying protein synthesis and degradation rates^24,70^) and protein perturbation techniques that lead to more precise and efficient manipulation of protein levels and activities; this would more clearly allow for investigating the association between change in transport upon knockdown and its original correlation to transport score from the single-cell data. In addition, subcellular proteomics may be subject to technical variability incurred during sample-preparation and nuclear isolation. To mitigate this effect, we utilized four biological replicates for single-nuclei and six biological replicates for bulk samples, and validated their nuclear enrichment, Additional file 1: Fig. S1. An additional consideration with nuclear isolation is that non-ionic detergents may permeabilize nuclear membranes, which we aimed to quantify, but the results for which were inconclusive, Additional file 1: Fig. S2. Similar future work would benefit from improved knockdown efficiency and more-stable and gentle nuclear isolation. Despite these potential experimental factors, this work identified and validated proteins associated with LPS-induced nucleocytoplasmic transport in macrophages using single-nucleus proteomics.

## Conclusions

This work demonstrates an approach which uses (sub-)single cell proteomics to infer functional regulators of complex biological processes, such as LPS-induced nucleocytoplasmic transport. We expect strategies such as this to aid in the functional interpretation of single-cell proteomics data and support the inference of novel functional regulators in diverse contexts.

## Methods

### Cell culture

THP-1 monocytes (TIB-202, ATCC) were cultured in RPMI-1640 Medium (R8758, Sigma-Aldrich) supplemented with 10% fetal bovine serum (10439016, Gibco) and 1% penicillin-streptomycin (15140122, Gibco) and grown at 37*^◦^*C and 5% CO_2_. Cells were transferred to 100×15mm Nunclon dishes (150350, Thermo Scientific) at a density of approximately 200,000 cells/mL and volume of 10 mL. Differentiation proceeded following an established protocol by Lund *et al.*^71^; THP-1 cells were incubated in the presence of 25 nM phorbol 12-myristate 13-acetate (PMA) for 48 hours, then cells were detached with accutase, mixed to prevent any batch-effects resulting from differentiation occurring in isolated plates, washed 2-times with phosphate-buffered saline (PBS), centrifuging at 300g for 5 min to pellet for washes. The cells were then allowed to incubate in growth media with-out PMA for 24 hours, and in this time reattach to fresh 100×15mm Nunclon dishes. The resulting M0 macrophages were either left not-treated (NT), or treated with 1 *µ*g/mL lipopolysaccharides from *Escherichia coli* O111:B4 (L4391, MilliporeSigma) for 10 minutes, 30 minutes, or 60 minutes. THP-1 cells used in these experiments did not undergo additional cell-line authentication, nor were they tested for mycoplasma contamination.

### Protein knockdowns

Proteins of interest were knocked down via reverse transfection of Silencer siRNAs (Ambion) and Silencer Select siRNAs (Ambion) into THP-1 monocytes using methodology adapted from manufacturer instructions for RNAi transfection. Silencer siRNAs were transfected in 12-well Nunclon plates and Silencer Select siRNAs were transfected in individual 60 mm dishes. For Silencer siRNAs, 10 pmol of siRNA was complexed with 3 *µL* of RNAiMAX. For Silencer Select siRNAs, 55 pmol of siRNA was complexed with 17 *µL* of RNAiMAX (Invitrogen) in 552 *µL* of serum-free RPMI (Sigma) in a 60 mm Nunclon dish (Thermo). siRNAs were complexed with the RNAiMAX lipid reagent for 5 minutes at room temperature. THP-1 monocytes were added in RPMI supplemented with 10% fetal bovine serum (FBS) and 1% penicillin-streptomycin to reach a final density of 300,000 cells per mL. Samples were cultured at 37*°*C and 5% CO_2_ for 24 hours before adding PMA, at which point the same aforementioned cell culture process was followed.

### Nuclear isolation

Macrophages were detached from Nunclon dishes with accutase on ice for approximately 20 minutes. The cells were washed twice with 1x PBS to remove media, centrifuging at 300g for 5 min each time. The cell concentration was measured with a hemocytometer. 15% of cells were retained as whole-cells for downstream analysis by resuspending the whole-cell pellet in LC-MS-grade water to a concentration of 3,000 cells/*µL*. These whole-cells were then stored at −80°C. The remaining 85% of cells were resuspended in 1 mL of ice-cold lysis buffer (0.1% NP-40, 250 mM sucrose, 25 mM KCl, 5 mM MgCl_2_, 10 mM HEPES (pH = 7.4)) in a 2 mL Eppendorf tube, and left to lyse on ice for 20 min. At this point in the protocol, the cells are lysed but the nuclei are unlikely to be free from cellular debris and other organelles. To purify the nuclei, the 1 mL nuclei suspension was sheared through a 25G needle with even pressure, avoiding bubbling and foaming as much as possible. The shearing was repeated for a total of 5 times. Nuclei were visually inspected under a microscope with trypan blue staining to ensure sufficient shearing of subcellular organelles from nuclei.

After nuclei are sufficiently sheared, the suspension was centrifuged at 1000g for 8 min at 4 *°*C. The nuclear pellet was resuspended in 500 µL of 250 mM sucrose, 25 mM KCl, 5 mM MgCl_2_, 10 mM HEPES (pH = 7.4). The mixture was underlayed with 500 µL of 350 mM sucrose, 25 mM KCl, 5 mM MgCl_2_, 10 mM HEPES (pH = 7.4) to form a sucrose cushion, which was then centrifuged at 4,000g for 10 min (4 *°*C). The supernatant was removed, and the nuclear pellet was resuspended in 200 µL of LC-MS water, gently flicking to mix. 4 µL of nuclei suspension was mixed with 4 µL of trypan blue, then counted with a hemocytometer to find the final concentration. 150 µL of the bulk nuclei were stored at −80°C. The remaining 50 µL was diluted to a concentration of roughly 400 nuclei/*µ*L and used for single-nuclei sorting and sample preparation by nPOP^16,57^.

#### Fluorescent dextran flow cytometry

Nuclei and whole cells were prepared from THP1 macrophage-like cells as reported in the manuscript. Each was split into 4 aliquots of 100 *µ*L cell or nuclei suspensions, each was left untreated or treated with 1 mg/mL of each fluorescent dextran reagent with average masses of 4 kDa (Sigma Aldrich, 46944-100MG-F), 40 kDa (Sigma Aldrich, FD40S100MG), and 150 kDa (Sigma Aldrich, 46946-100MG-F) for 20 minutes on ice, washed with 1 mL PBS or nuclei buffer (20% iodixanol, 25 mM KCl, 5 mM MgCl2, 10 mM HEPES (pH = 7.4)), and then resuspended in 100 *µ*L of their respective buffers. The suspensions of whole cells and nuclei were then analyzed by flow cytometry using a Beckman Coulter CytoFLEX Flow Cytometer.

### Sample preparation for proteomic analysis

#### Bulk samples

Nuclei and whole-cell fractions, which were frozen at −80°C at a concentration of 3,000 cells/*µ*L, were heated to 90°C for 10 minutes as part of the mPOP protocol^72^. Trypsin Gold (V5280, Promega) was added to a final concentration of 20 ng/*µ*L, in addition to final concentrations of 100 mM triethylammonium bicarbonate (TEAB) and 0.25 U/*µ*L benzonase nuclease (E1014, Millipore Sigma). The cell lysates were digested at 37°C for 18 hours in a thermocycler. Samples were labeled with mTRAQ following manufacturer’s instructions. In short, the labels, which arrived suspended in isopropyl alcohol at a concentration of 0.05 U/*µ*L (where 1 unit labels 100 *µ*g), were added in a 1:2 proportion of label:peptide where the peptides were concentrated to 500 ng/*µ*L in 100 mM TEAB. The labeling reaction was left at room temperature for 2 hours, and then quenched with 0.25% hydroxylamine for 1 hour.

#### Creating a mixed species standard to benchmark SCoPE-DIA

Digested protein lysate from THP-1 macrophage nuclei were mixed with digested *S. cerevisiae* protein lysate (V7461, Promega) to generate three samples. The proportions of Sample A (Δ0):Sample B (Δ4) was 4:1 for *S. cerevisiae* and 1:1 for *H. sapiens*. The unlabeled versions of these samples were mixed in a 1:1 ratio to generate a carrier sample which was then labeled with mTRAQ Δ8.

Samples A and B were diluted such that the *H. sapiens* amount was 1 nucleus worth of protein (approximately 45 pg). The carrier sample was pooled with Samples A and B at 0x, 1x, 5x, 10x, 25x, and 50x concentrations to generate the SCoPE-DIA benchmarks.

#### Single nuclei

Single nuclei were prepared for proteomic analysis by nPOP as previously described by Leduc *et al.*^16,57^. In short, single-nuclei were sorted by CellenOne into 8 nL droplets of 100% DMSO on a fluorocarbon-coated glass slide to lyse. The lysed nuclei were digested with Trypsin Gold at a concentration of 120 ng/*µ*L and 5 mM HEPES (pH 8.5) and 200 mM TEAB buffer (pH 8.5). Peptides were then chemically labeled with mTRAQ Δ0 or Δ4. Single nuclei were pooled into sets of Δ0 and Δ4, and dispensed into individual wells of a 384 well-plate. The pooled samples in the 384 well-plate were then dried in a speed-vacuum then stored at −80°C until use. Before sample loading for LC-MS/MS, a 25x or 50x nuclei carrier labeled with mTRAQ Δ8 was pooled into each set to create the final Δ0, Δ4, and Δ8 SCoPE-DIA set. Four single-nucleus biological replicates were prepared and acquired, for a total of 3412 single nuclei.

### Data acquisition

All single nuclei and bulk samples were resuspended in 0.01% n-Dodecyl *β*-d-maltoside (DDM)^73^ with 0.1% formic acid. Sample pickup occurred out of a 384 well-plate for single-nucleus samples, or from glass vials for bulk samples. Peptides were separated for LC-MS acquisition using a Neo Vanquish UHPLC and 25 cm × 75 *µm* IonOpticks Aurora Series UHPLC columns (AUR2-25075C18A) at a flow-rate of 200 nL/min. LC was run with the following settings: Direct Injection, nano/cap flow, maximum pressure = 1500 bar, maximum pressure change = 1000 bar/min.

Sample loading had the following settings: 1*µ*L injection with 1.2 *µ*L loading volume, “Pressure Control” mode with maximum pressure set to 1450 bar, and fast loading enabled. Wash and equilibration settings: “Pressure Control” mode with maximum pressure set to 1450 bar, equilibration factor = 4.0, and fast equilibration was enabled.

The LC gradient proceeded as follows: Ramp from 2.5%B to 6.5%B over 0.2 minutes, to 11.5%B over 0.9 minutes, to 21.0%B over 3.1 minutes, to 31.5%B over 6.2 minutes, to 40%B over 2.8 minutes, to 55%B over 1.7 minutes, to 95%B over 0.65 minutes and then hold at 95%B for 4 minutes. This method allows for approximately 15 minutes of active chromatography, at a through-put of approximately 45 runs/day (90 single nuclei/day with SCoPE-DIA or 135 bulk samples/day with plexDIA) after accounting for sample loading and column equilibration overheads.

All data were acquired on a Bruker timsTOF SCP using captive spray, dia-PASEF scan mode^74,75^, and positive polarity. The duty cycle consisted of 8 total PASEF frames, 26 Th MS2 windows with 1 Th overlaps. To improve temporal sampling of the MS1-elution profiles which are used for quantification, an MS1 scan was taken every 2 PASEF frames, for a total of 4 MS1 scans per duty cycle. MS2 scan range: 300 m/z-1000 m/z, MS1 scan range: 100 m/z-1700 m/z, 1/K0 start: 0.64, 1/K0 end: 1.20, ramp and accumulation times: 100 ms. The estimated duty cycle time is 1.28 seconds. Collision energy settings were 20 eV at 1/K0 = 0.60 and 59 eV at 1/K0 = 1.60. Collision RF was set to 2000 Vpp.

### Raw data searching with DIA-NN

Empirical spectral libraries of mTRAQ-labeled nuclei and whole-cells were generated from di-aPASEF runs on the timsTOF SCP, searching with a DIA-NN *in silico*-predicted Swiss-Prot *H. sapiens* FASTA (canonical & isoform). Similarly, a combined *H. sapiens* and *S. cerevisiae* library was generated for searching SCoPE-DIA benchmarks.

DIA-NN (version 1.8.1) was used to search raw data as previously implemented^25,76^. The following search settings were used: {–fixed-mod mTRAQ 140.0949630177, nK}, {–channels mTRAQ, 0, nK, 0:0; mTRAQ, 4, nK, 4.0070994:4.0070994; mTRAQ, 8, nK, 8.0141988132:8.0141988132}, {–original-mods}, {–peak-translation}, {–report-lib-info}, {–qvalue 0.01}, {–mass-acc-ms1 5.0}, {–mass-acc 15.0}, {–mass-acc-quant 5.0}, {–reanalyse}, {–rt-profiling}, and {–peak-height}.

### Computational data analysis

#### Nuclear enrichment

To assess nuclear enrichment, the first two bulk biological replicates were acquired in a plexDIA set of whole-cell and nuclear fractions. The fractions were subset for histones, which should only be present in the nucleus, and both fractions were normalized by a scalar such that histone proteins between whole-cells and nuclear fractions were in a 1:1 ratio. Ratios of MaxLFQ protein abundances for protein markers from The Human Protein Atlas for endoplasmic reticulum (ER), nuclear compartments, mitochondria, plasma membrane, cytosol, and Golgi were calculated between nuclear and whole-cell fractions to assess nuclear protein enrichment^77–79^. These ratios were averaged per-protein across replicates.

#### Differential protein abundance analysis

Precursor quantities acquired by plexDIA and analyzed by DIA-NN were corrected for isotopic envelope carryover, as previously performed^25^. These precursor quantities were filtered for Lib.PG.Q.Value *<* 0.01 and Q.Value *<* 0.01 for bulk data from all six biological replicates and their corresponding nested technical replicates. The bulk data used to generate weights for single-nuclei analyses were taken from cell-batches that corresponded to those single nuclei, in this case, biological replicates 1, 2, 3, and 6; the bulk data used for single-nucleus weighting were further filtered for Translated Q-value and Channel Q-value *<* 0.05.

Differential protein abundance was calculated through the MS-EmpiRe workflow^42^ for each biological replicate. Precursors used for analysis were required to be quantified in at least two replicates per condition. Data normalization was performed using operations from the MS-EmpiRe workflow, as described by Ammar *et al.*^42^. Differential abundance analysis collapsed precursors to gene-level annotations; this produced p-values calculated after outlier correction for each protein for each biological replicate; Stouffer’s method was applied to collapse multiple p-values from several biological replicates to a single combined p-value. Briefly, this involved converting p-values for each protein from each biological replicate to z-scores, multiplying by the sign of the fold-change, summing the signed z-scores, then dividing by the square root of the number of observations for that protein, and finally converting that z-score back to a single p-value for each protein. The resulting p-values were corrected for multiple hypothesis testing using the Benjamini-Hochberg (BH) correction^80^.

#### Protein set enrichment analysis

Protein set enrichment analysis was performed using the g:GOSt R package^81,82^. An ordered vector of gene names was used as the input. Results were filtered at 1% FDR, and the relative abundance for the Gene Ontology terms was computed across all samples from the intersect of protein abundances to represent the relative abundance of a given Gene Ontology term. Calculating the relative abundance of a Gene Ontology term from the subset of intersected proteins allowed for fairer comparisons of enrichment between samples. Specifically regarding the analysis of whole-cell bulk knockdowns shown in Additional file 1: Fig. S14, in some cases this was performed on data in which NT and 60 minute LPS-treated cell-lysates were combined. These samples were experimentally combined for ease of labor and due to the minimal impact of 60 minute LPS-treatment on the whole-cell proteome.

#### Evaluating relationship of protein mass and transport

Protein masses were downloaded from UniProt^83^, and matched to proteins which we found significantly increased or decreased in response to LPS (Q-value *<* 0.05) in nuclear fractions. The absolute value of the log_2_ fold-change was plotted with the mass of the protein to statistically evaluate the relationship between protein mass and the amount of transport that occurred as a result of LPS treatment. Because the masses are purely based on gene-encoded sequences, this analysis does not account for changes in mass as the result of proteolytic cleavage or the transport of protein complexes.

Additionally, the dynamics of protein transport were interpolated from the four data points (NT, 10 min, 30 min, and 60 min LPS) using a 3rd degree polynomial for each of the proteins that was found to significantly change in at least one of the conditions relative to NT. A total of 300 time points were predicted from the fit, ranging from 0 to 60 minutes. The proteins were arranged from smallest to largest and a moving median of the absolute value of the log_2_ fold-change from NT was calculated for each time point from that protein’s adjacent 40 smaller and 40 larger proteins. In this way, biological differences between proteins could be averaged to generally assess the influence of protein mass on transport kinetics. For each protein’s averaged kinetics, the time when half of the total absolute value log_2_ fold-change had occurred was marked. This was plotted against the protein mass and a Spearman correlation was calculated.

#### Benchmarking SCoPE-DIA quantification

To benchmark how different levels of carrier affect protein-level quantitation of SCoPE-DIA as shown in Additional file 1: Fig. S6f, precursors were intersected across all carrier levels, and thus proteins as well; precursor quantities were collapsed to protein-level quantities using MaxLFQ, and then ratio values between Δ0 and Δ4 were computed, normalizing *H. sapiens* to a be mediancentered at a 1:1 ratio to account for systematic differences in loading amounts, if any exist. To compare quantitation at different carrier levels for all proteins, as shown in Additional file 1: Fig. S6f, the same process was performed without intersecting.

In regards to compression filtering, acquiring single cell (or nucleus) data with a constant, known amount of carrier is useful to identify precursors which have systematically interfered quantification. Precursors which have systematically compressed quantification across all samples relative to the expected carrier amount, are unlikely to be well-quantified and can be removed. To identify which precursors to remove, a ratio for each precursor from the single-nucleus channels to its respective carrier channel was calculated. Precursors with median ratios *>* 6-fold greater than the theoretical ratio (e.g. 1:50 is expected for a 50x carrier) were considered to be systematically poorly quantified and were excluded from downstream analyses.

#### Single-nucleus quality control

To identify which single nuclei were successfully prepared and acquired, precursors were filtered for Channel and Translated Q-values *<* 0.1 and the number of remaining high quality precursors was counted. Single nuclei with *≥* 50 high quality precursors were retained for downstream analysis as this threshold was sufficiently separable from negative controls. Precursors were required to be quantified in *≥* 1% of single nuclei, and have *<* 6x ratio compression to be retained for downstream analysis.

Single nuclei were further filtered to ensure sufficient depletion of non-nuclear proteins. To quantify the nuclear purity of each single-nucleus, the nuclear proteomes were scaled to a reference bulk nuclear fraction, and compared to a ratio of whole-cell to nuclear fraction. A 3rd degree polynomial was used to fit a curve for each single-nucleus, and an absolute value of the area under the curve (AUC) was computed. This value was used to identify single-nuclei which were insufficiently pure. Only single nuclei with AUCs *<* 5 were used in downstream analyses.

#### Single-nucleus protein quantification, imputation, and batch correction

Precursor abundances from single nuclei were normalized to their respective carrier channel, which serves as a reference similar to the isobaric SCoPE-MS workflow^12,27^. For each precursor across all carrier samples, the mean abundance was calculated and used to scale the data of the single nuclei, to be compatible for MaxLFQ protein quantification^77^. The resulting protein-level data of each nucleus was normalized by its respective median protein abundance to account for differences in absolute abundances, then each protein was normalized by the mean abundance across single nuclei. The log_2_-transformed data was imputed using a weighted *k*-nearest neighbors (kNN) approach (*k*=5), where the weights were proportional to the similarity to its nearest neighbors. Finally, the data was batch-corrected using ComBat^84,85^ to account for mTRAQ label-biases, LC-batches, and biological replicates.

#### Weighted principal component analysis (PCA)

The variance of single-nucleus protein data was weighted based on protein correlations, bulk differential abundance, and bulk continuity of spatiotemporal trends. Effectively these weights assign higher variance based on: 1) how correlated a protein’s abundance is to other proteins abundances, 2) how differentially abundant the protein was in the bulk data, and 3) how continuous the trend was in its spatiotemporal response to LPS.

Specifically, the bulk differential abundance data computed from MS-EmpiRe^42^ analysis had p-values which were converted to signed Z-scores. Using Stouffer’s method they were combined for all the bulk biological replicates that correspond to the single-nucleus replicates, in this case, biological replicates 1, 2, 3, and 6. The resulting Z-score for each protein was squared to generate non-negative weights, then used to weight the variance of that protein in the single-nucleus data. Additionally, the inverse of the ranked version of the von Neumann ratio (RVN) calculated from bulk data was used to weight the variance of each respective protein in the single-nucleus data. Eigendecomposition was then performed on the cell-cell correlation matrix.

#### Computing single-nucleus transport scores

Directly measuring protein transport in a cell using LC-MS/MS was not possible with this experimental design as we did not measure the same nucleus before and after LPS stimulation. However, we derived a ‘transport score’ metric to serve as a substitute which serves to quantify the deviation of single LPS-treated nuclei from the population of single NT nuclei. The transport score accounts for pre-existing variability in the not-treated (NT) population of single nuclei for each protein by converting protein abundances for each LPS-treated nucleus to a Z-score in reference to the NT-population. For each nucleus, the resulting vector of Z-scores is weighted based on the signed-Z-scores of the differential protein abundances in the bulk data. Therefore, a protein’s contribution to the transport score is weighted by its significance in the bulk data, where more differential proteins carry greater weights. The mean component of the resulting vector of each nucleus is taken as that nucleus’ transport score.

#### Computing protein overlap coefficients between populations of single nuclei

The overlap between the distributions of relative protein abundances between populations of single nuclei was computed to quantify the commonality of protein abundances pre-stimulation and post-LPS stimulation. This overlap coefficient was computed by the overlap function of bayestestR^86^. It is computed by modeling the distribution of relative protein abundances as densities for two populations and then computing the proportion that is shared. For example, a protein that has high-overlap between abundances in NT and LPS-treated populations, is one that is minimally transported relative to its natural variability.

#### Quantifying nuclear pore complex abundances and dependence on mass

To quantify nuclear pore complex abundances in each single-nucleus, relative abundances of nucleoporins were converted to Z-scores and combined for all nucleoporins using Stouffer’s method. Mass dependence as a function of NPC abundance was computed for each LPS-treated single-nucleus based on the slopes (protein Z-scores vs log_10_-transformed protein mass) of differentially abundant proteins in the single-nucleus data.

#### Investigating the association between nucleoporin half-lives and their correlations to transport score

Half-lives were derived from SILAC turnover data of primary monocytes, NK cells, neurons, hepatocytes, and B-cells from Mathieson *et al.*^62^. Specifically, the half-lives of all replicates for each cell-type was collapsed to the mean, then the mean of all cell-types was collapsed to a final value, which was used as the half-life in our analysis.

#### Visualizing the nuclear pore complex

The predicted structure of the human nuclear pore complex (PDB Entry: 7r5k)^61^ from Mosalaganti *et al.* was used visualize the NPC. The nucleoporins were colored by their respective correlations to transport score using ChimeraX (v1.6.1)^87^.

#### Quantifying the effect of protein knockdowns on transport

To quantify how each protein knockdown affected LPS-induced nucleocytoplasmic transport, each biological replicate included a matched negative (non-targeting) siRNA control to account for how much transport would have occurred without the knockdown. A slope was calculated from the 100 most differentially abundant proteins in the original (no knockdown) bulk data to the data for each knockdown. The fold-changes for proteins used for computing these slopes were weighted by their associated Z-score significance of differential abundances from the original bulk data; this serves to weight each protein’s contribution to the slope-calculation based on its significance. The difference in slopes between the negative control (non-targeting) siRNA and the knockdown was computed. This difference in slope is interpreted as the extent to which nucleocytoplasmic transport changed as a result of the knockdown. Additionally, the associated standard error of each slope was used between the negative control and experimental knockdown to derive a Z-score for the significance of the difference in transport. Significance from additional biological replicates were combined using Stouffer’s method to arrive at a cumulative Z-score. To assess the significance of the effect on transport of individual knockdowns, this Z-score was converted to a p-value and adjusted for multiple testing hypotheses using a BH-correction. To summarize, this approach statistically evaluates the difference in global LPS-induced transport (quantified as slopes) between cell batches that were treated or not treated with a specific siRNA.

#### Calculating significance of correlation to transport score

To derive a metric for the significance of protein correlations to the transport score, a rank-based normalization approach (Rankit) was used to transform the correlations into a standard normal distribution. The resulting Z-scores were combined using Stouffer’s method to arrive at a final Z-score for each protein, as shown on the y-axis of Fig. 4c. This approach was chosen as an alternative to a traditional Z-score operation given that the distribution of correlations to transport scores were non-Gaussian and have a skewed tail.

## Declarations

### Ethics approval and consent to participate

Not applicable.

### Consent for publication

Not applicable.

### Availability of data and materials

All data are reported and deposited in accordance with the community guidelines^22^. Raw data, spectral libraries, FASTAs, and DIA-NN search outputs are available at MassIVE: MSV000094829^88^. Code, processed data, supporting files, supplementary figures, and interactive supplementary analyses (e.g. volcano plots and time-series data) are available at github.com/SlavovLab/Single-nucleus-proteomics^89^ and are archived at Zenodo (doi.org/10.5281/zenodo.24144392)^90^. The source code is released under the MIT license.

### Authors’ contributions

**Experimental design**: J.D., T.J., and N.S.

**Data analysis**: J.D.

**LC-MS/MS**: J.D. and T.J.

**Sample preparation**: J.D., T.J., A.L., L.K., S.K., and M.R.

**Raising funding**: N.S.

**Supervision**: N.S.

**Initial draft**: J.D., T.J., and N.S.

All authors read and approved the final manuscript.

## Acknowledgements

We thank H. Specht, R. Huffman, A. Petelski, D. Perlman, and E. Emmott for discussions per-taining to experimental methodology for single-nucleus proteomics, M. Rout and T. Eeuwen for thoughtful discussions regarding nuclear pore complexes, and I. Zanoni and M. Di Gioia for discussions regarding LPS-response in macrophages and early experimental work. Flow cytometry was performed in the Institute for Chemical Imaging of Living Systems (CILS) at Northeastern University.

## Peer review information

Veronique van den Berghe was the primary editor of this article and managed its editorial process and peer review in collaboration with the rest of the editorial team. The peer-review history is available in the online version of this article.

## Additional files

**Additional file 1** (.pdf): Supplementary Figs. S1–S14, with legends. Supplementary figures on bulk nuclear enrichment, fluorescent dextran experiments, time-series continuity, mass dependence of transport, and DNA repair proteins; benchmarking of SCoPE-DIA and single-nucleus quality control; calculation of the transport score and protein set enrichment of correlations to transport score; validation of siRNA knockdowns, per-replicate changes in transport, robustness to filtering, and whole-cell protein set enrichment of knockdowns.

**Additional file 2** (.xlsx): Supplementary Tables S1–S17: sample annotations, protein abundances, differential abundance, protein set enrichment analyses, correlations to transport score, nucleo-porin correlations, and siRNA knockdown results.

## References

1. Macosko, E. Z. et al. Highly parallel genome-wide expression profiling of individual cells using nanoliter droplets. Cell 161, 1202–1214 (2015).

2. Slavov, N. Scaling Up Single-Cell Proteomics. Molecular & Cellular Proteomics 21, 100179 (2022).

3. Quake, S. R. A decade of molecular cell atlases.

4. Symmons, O. & Raj, A. Whats Luck Got to Do with It: Single Cells, Multiple Fates, and Biological Nondeterminism. Molecular cell 62, 788–802 (2016).

5. Slavov, N. Unpicking the proteome in single cells. Science 367, 512–513 (2020).

6. Weinreb, C., Wolock, S., Tusi, B. K., Socolovsky, M. & Klein, A. M. Fundamental limits on dynamic inference from single-cell snapshots. Proceedings of the National Academy of Sciences 115, E2467–E2476 (2018).

7. Leduc, A., Harens, H. & Slavov, N. Modeling and interpretation of single-cell proteogenomic data. arXiv. eprint: 2308.07465 (2023).

8. Munsky, B., Neuert, G. & van Oudenaarden, A. Using gene expression noise to understand gene regulation. Science 336, 183–187 (2012).

9. Slavov, N. Learning from natural variation across the proteomes of single cells. PLOS Biology 20, 1–4 (2022).

10. Kelly, R. T. Single-Cell Proteomics: Progress and Prospects. Molecular & Cellular Pro-teomics (2020).

11. Slavov, N. Single-cell protein analysis by mass spectrometry. Current Opinion in Chemical Biology 60, 1–9 (2020).

12. Specht, H. et al. Single-cell proteomic and transcriptomic analysis of macrophage heterogeneity using SCoPE2. Genome Biology 22, 50 (2021).

13. Cong, Y. et al. Ultrasensitive single-cell proteomics workflow identifies *>* 1000 protein groups per mammalian cell. Chemical Science (2020).

14. Huffman, R. G. et al. Prioritized mass spectrometry increases the depth, sensitivity and data completeness of single-cell proteomics. Nature Methods 20, 714–722 (2023).

15. Thielert, M. et al. Robust dimethyl-based multiplex-DIA doubles single-cell proteome depth via a reference channel. Mol. Syst. Biol. 19, e11503 (2023).

16. Leduc, A., Huffman, R. G., Cantlon, J., Khan, S. & Slavov, N. Exploring functional protein covariation across single cells using nPOP. Genome Biology 23, 261 (2022).

17. Orsburn, B. C., Yuan, Y. & Bumpus, N. N. Insights into protein post-translational modification landscapes of individual human cells by trapped ion mobility time-of-flight mass spectrometry. Nature Communications 13, 7246 (2022).

18. Papalexi, E. & Satija, R. Single-cell RNA sequencing to explore immune cell heterogeneity. Nature Reviews Immunology 18, 35–45 (2018).

19. Peters-Clarke, T. M. et al. Boosting the Sensitivity of Quantitative Single-Cell Proteomics with Infrared-Tandem Mass Tags. J. Proteome Res. (2024).

20. Peters-Clarke, T. M., Coon, J. J. & Riley, N. M. Instrumentation at the Leading Edge of Proteomics. Analytical Chemistry.

21. Slavov, N. Driving Single Cell Proteomics Forward with Innovation. Journal of Proteome Research 20, 4915–4918 (2021).

22. Gatto, L. et al. Initial recommendations for performing, benchmarking, and reporting single-cell proteomics experiments. Nat. Methods 20, 375–386 (2023).

23. Slavov, N. Single-Cell Proteomic Technologies: Tools in the quest for principles. Annu. Rev. Biophys. 55, 253–275 (2026).

24. Slavov, N. Developing Single-Cell Proteomics to Open New Perspectives on Biology. Journal of the American Society for Mass Spectrometry (2026).

25. Derks, J. et al. Increasing the throughput of sensitive proteomics by plexDIA. Nat. Biotechnol. (2022).

26. Derks, J. & Slavov, N. Strategies for increasing the depth and throughput of protein analysis by plexDIA. Journal of Proteome Research 22, 697–705 (2023).

27. Budnik, B., Levy, E., Harmange, G. & Slavov, N. SCoPE-MS: mass spectrometry of single mammalian cells quantifies proteome heterogeneity during cell differentiation. Genome Biology 19, 161 (2018).

28. Specht, H. & Slavov, N. Optimizing Accuracy and Depth of Protein Quantification in Experiments Using Isobaric Carriers. Journal of Proteome Research 20, 880–887 (2021).

29. Welter, A. S., et al. Combining data independent acquisition with spike-in SILAC (DIA-SiS) improves proteome coverage and quantification. bioRxiv (2024).

30. Van Bentum, M. et al. Spike-in enhanced phosphoproteomics uncovers synergistic signaling responses to MEK inhibition in colon cancer cells 2024.

31. Khoury, L., et al. Single-cell proteomics reveals cell-type-specific functional coordination in PBMCs. bioRxiv (2026).

32. Tay, S. et al. Single-cell NF-*κ*B dynamics reveal digital activation and analogue information processing. Nature 466, 267–271 (2010).

33. Son, M. et al. Spatiotemporal NF-*κ*B dynamics encodes the position, amplitude, and duration of local immune inputs. Sci Adv 8, eabn6240 (2022).

34. Levy, E. & Slavov, N. Single cell protein analysis for systems biology. Essays In Biochemistry 62 (4 2018).

35. Wang, A. G., Son, M., Kenna, E., Thom, N. & Tay, S. NF-*κ*B memory coordinates transcriptional responses to dynamic inflammatory stimuli. Cell reports 40 (2022).

36. Zhang, Q. et al. NF-*κ*B dynamics discriminate between TNF doses in single cells. Cell systems 5, 638–645 (2017).

37. Lee, R. E. C., Walker, S. R., Savery, K., Frank, D. A. & Gaudet, S. Fold change of nuclear NF-*κ*B determines TNF-induced transcription in single cells. Molecular Cell 53, 867–879 (2014).

38. Noursadeghi, M. et al. Quantitative imaging assay for NF-kappaB nuclear translocation in primary human macrophages. Journal of Immunological Methods 329, 194–200 (2008).

39. Sakai, N. et al. Direct Visualization of the Translocation of the *γ*-Subspecies of Protein Kinase C in Living Cells Using Fusion Proteins with Green Fluorescent Protein. Journal of Cell Biology 139, 1465–1476 (1997).

40. Wang, Y., Shyy, J. Y.-J. & Chien, S. Fluorescence Proteins, Live-Cell Imaging, and Mechanobiology: Seeing Is Believing. Annual Review of Biomedical Engineering 10, 1–38 (2008).

41. Krishnaswami, S. R. et al. Using single nuclei for RNA-seq to capture the transcriptome of postmortem neurons. Nature Protocols 11, 499–524 (2016).

42. Ammar, C., Gruber, M., Csaba, G. & Zimmer, R. MS-EmpiRe Utilizes Peptide-level Noise Distributions for Ultra-sensitive Detection of Differentially Expressed Proteins[S]. Molecular & Cellular Proteomics 18, 1880–1892 (2019).

43. Bartels, R. The Rank Version of von Neumann’s Ratio Test for Randomness. Journal of the American Statistical Association 77, 40–46 (1982).

44. Ghavami, A., van der Giessen, E. & Onck, P. R. Energetics of Transport through the Nuclear Pore Complex. PLoS ONE 11, e0148876 (2016).

45. Wing, C. E., Fung, H. Y. J. & Chook, Y. M. Karyopherin-mediated nucleocytoplasmic transport. Nature Reviews Molecular Cell Biology 23, 307–328 (2022).

46. Keminer, O. & Peters, R. Permeability of single nuclear pores. Biophysical Journal 77, 217–228 (1999).

47. Paci, G., Zheng, T., Caria, J., Zilman, A. & Lemke, E. A. Molecular determinants of large cargo transport into the nucleus. eLife 9, e55963 (2020).

48. Timney, B. L. et al. Simple rules for passive diffusion through the nuclear pore complex. Journal of Cell Biology 215, 57–76 (2016).

49. Beinke, S. & Ley, S. C. Functions of NF-*κ*B1 and NF-*κ*B2 in immune cell biology. Biochemical Journal 382, 393–409 (2004).

50. Liu, T., Zhang, L., Joo, D. & Sun, S.-C. NF-*κ*B signaling in inflammation. Signal Transduction and Targeted Therapy 2, 1–9 (2017).

51. Mulvey, C. M. et al. Spatiotemporal proteomic profiling of the pro-inflammatory response to lipopolysaccharide in the THP-1 human leukaemia cell line. Nature Communications 12, 5773 (2021).

52. Zhou, H.-R., Islam, Z. & Pestka, J. J. Kinetics of lipopolysaccharide-induced transcription factor activation/inactivation and relation to proinflammatory gene expression in the murine spleen. Toxicology and Applied Pharmacology 187, 147–161 (2003).

53. Ochi, T. et al. PAXX, a paralog of XRCC4 and XLF, interacts with Ku to promote DNA double-strand break repair. Science (New York, N.Y.) 347, 185–188 (2015).

54. Xing, M. et al. Interactome analysis identifies a new paralogue of XRCC4 in non-homologous end joining DNA repair pathway. Nature Communications 6, 6233 (2015).

55. Liu, X., Shao, Z., Jiang, W., Lee, B. J. & Zha, S. PAXX promotes KU accumulation at DNA breaks and is essential for end-joining in XLF-deficient mice. Nature Communications 8, 13816 (2017).

56. McCloskey, A., Ibarra, A. & Hetzer, M. W. Tpr regulates the total number of nuclear pore complexes per cell nucleus. Genes & Development 32, 1321–1331 (2018).

57. Leduc, A., Khoury, L., Cantlon, J., Khan, S. & Slavov, N. Massively parallel sample preparation for multiplexed single-cell proteomics using nPOP. Nature Protocols, 1–27 (2024).

58. Petelski, A. A. et al. Multiplexed single-cell proteomics using SCoPE2. Nature Protocols 16, 5398–5425 (2021).

59. Van Bergen, N. J. et al. Pathogenic variants in nucleoporin TPR (translocated promoter region, nuclear basket protein) cause severe intellectual disability in humans. Human Molecular Genetics 31, 362–375 (2022).

60. Han, L. et al. Changes in nuclear pore numbers control nuclear import and stress response of mouse hearts. Developmental Cell 57, 2397–2411.e9 (2022).

61. Mosalaganti, S. et al. AI-based structure prediction empowers integrative structural analysis of human nuclear pores. Science 376, eabm9506 (2022).

62. Mathieson, T. et al. Systematic analysis of protein turnover in primary cells. Nature Communications 9, 689 (2018).

63. Savas, J. N., Toyama, B. H., Xu, T., Yates, J. R. & Hetzer, M. W. Extremely Long-lived Nuclear Pore Proteins in the Rat Brain. Science (New York, N.y.) 335, 942 (2012).

64. Raman, N., Weir, E. & Müller, S. The AAA ATPase MDN1 Acts as a SUMO-Targeted Regulator in Mammalian Pre-ribosome Remodeling. Molecular Cell 64, 607–615 (2016).

65. Schink, S., Ammar, C., Chang, Y.-F., Zimmer, R. & Basan, M. Analysis of proteome adaptation reveals a key role of the bacterial envelope in starvation survival. Molecular Systems Biology 18, e11160 (2022).

66. Ori, A. et al. Cell type-specific nuclear pores: a case in point for context-dependent stoichiometry of molecular machines. Molecular systems biology 9, 648 (2013).

67. Wang, C. et al. High-throughput label-free single-cell proteomics enabled by multicolumn NanoLC. Nature Communications (2026).

68. Specht, H., et al. PSMtags improve peptide sequencing and throughput in sensitive proteomics. bioRxiv (2025).

69. Derks, J., et al. Increasing mass spectrometry throughput using time-encoded sample multiplexing. bioRxiv (2025).

70. Leduc, A., Shipkovenska, G., Xu, Y., Franks, A. & Slavov, N. Principles of protein abundance regulation across single cells in a mammalian tissue. bioRxiv (2025).

71. Lund, M. E., To, J., O’Brien, B. A. & Donnelly, S. The choice of phorbol 12-myristate 13-acetate differentiation protocol influences the response of THP-1 macrophages to a pro-inflammatory stimulus. Journal of Immunological Methods 430, 64–70 (2016).

72. Specht, H. et al. Automated sample preparation for high-throughput single-cell proteomics 2018.

73. Dou, M. et al. Automated Nanoflow Two-Dimensional Reversed-Phase Liquid Chromatography System Enables In-Depth Proteome and Phosphoproteome Profiling of Nanoscale Samples. Analytical Chemistry 91, 9707–9715 (2019).

74. Meier, F. et al. diaPASEF: parallel accumulation–serial fragmentation combined with data-independent acquisition. Nature Methods 17, 1229–1236 (2020).

75. Demichev, V. et al. dia-PASEF data analysis using FragPipe and DIA-NN for deep proteomics of low sample amounts. Nature Communications 13, 3944 (2022).

76. Demichev, V., Messner, C. B., Vernardis, S. I., Lilley, K. S. & Ralser, M. DIA-NN: neural networks and interference correction enable deep proteome coverage in high throughput. Nature methods 17, 41–44 (2020).

77. Cox, J. et al. Accurate Proteome-wide Label-free Quantification by Delayed Normalization and Maximal Peptide Ratio Extraction, Termed MaxLFQ. Molecular & Cellular Proteomics: MCP 13, 2513–2526 (2014).

78. Thul, P. J. & Lindskog, C. The human protein atlas: A spatial map of the human proteome. Protein Science : A Publication of the Protein Society 27, 233–244 (2018).

79. Pontén, F., Jirstrém, K. & Uhlen, M. The Human Protein Atlas—a tool for pathology. The Journal of Pathology 216, 387–393 (2008).

80. Benjamini, Y. & Hochberg, Y. Controlling the False Discovery Rate: A Practical and Powerful Approach to Multiple Testing. Journal of the Royal Statistical Society. Series B (Methodological) 57, 289–300 (1995).

81. Reimand, J., Kull, M., Peterson, H., Hansen, J. & Vilo, J. g:Profiler—a web-based toolset for functional profiling of gene lists from large-scale experiments. Nucleic Acids Research 35, W193–W200 (2007).

82. Kolberg, L. et al. g:Profiler—interoperable web service for functional enrichment analysis and gene identifier mapping (2023 update). Nucleic Acids Research 51, W207–W212 (2023).

83. The UniProt Consortium. UniProt: the Universal Protein Knowledgebase in 2023. Nucleic Acids Research 51, D523–D531 (2023).

84. Johnson, W. E., Li, C. & Rabinovic, A. Adjusting batch effects in microarray expression data using empirical Bayes methods. Biostatistics (Oxford, England) 8, 118–127 (2007).

85. Leek, J. T., Johnson, W. E., Parker, H. S., Jaffe, A. E. & Storey, J. D. The sva package for removing batch effects and other unwanted variation in high-throughput experiments. Bioinformatics 28, 882–883 (2012).

86. Makowski, D., Ben-Shachar, M. S. & Lüdecke, D. bayestestR: Describing Effects and their Uncertainty, Existence and Significance within the Bayesian Framework. Journal of Open Source Software 4, 1541 (2019).

87. Goddard, T. D. et al. UCSF ChimeraX: Meeting modern challenges in visualization and analysis. Protein Science 27, 14–25 (2018).

88. Derks, J., Jonson, T., Leduc, A., Rafiee, M.-R., Khan, S., Khoury, L. & Slavov, N. Single-nucleus proteomics identifies regulators of protein transport. Datasets. MassIVE. 10.25345/C5959CK32 (2024).

89. Derks, J., Jonson, T., Leduc, A., Rafiee, M.-R., Khan, S., Khoury, L. & Slavov, N. Single-nucleus-proteomics. GitHub. https://github.com/SlavovLab/Single-nucleus-proteomics (2024).

90. Derks, J., Jonson, T., Leduc, A., Rafiee, M.-R., Khan, S., Khoury, L. & Slavov, N. Single-nucleus-proteomics. Zenodo. 10.5281/zenodo.18773852 (2026).

